# Scene-selective brain regions respond to embedded objects of a scene

**DOI:** 10.1101/2022.06.22.497191

**Authors:** Elissa M. Aminoff, Tess Durham

## Abstract

Objects are fundamental to scene understanding. Scenes are defined by embedded objects and how we interact with them. Paradoxically, scene processing in the brain is typically discussed *in contrast* to object processing. Using the BOLD5000 dataset (Chang et al., 2019), we examined whether objects within a scene predicted the neural representation of scenes, as measured by fMRI in humans. Stimuli included 1,179 unique scenes across 18 semantic categories. Object composition of scenes were compared across scene exemplars in different semantic categories, and separately, in exemplars of the same category. Neural representations in scene- and object-preferring brain regions were significantly related to which objects were in a scene, with the effect at times stronger in the scene-preferring regions. The object model accounted for more variance when comparing scenes within the same semantic category to scenes from different categories. Thus, the functional role of scene-preferring regions should include the processing of objects. This suggests visual processing regions may be better characterized with respect to which processes are engaged when interacting with the stimulus category, such as processing groups of objects in scenes, or processing a single object in our foreground, rather than the stimulus category itself.

---

The nature of scene understanding is fundamentally tied to the objects that comprise scenes. The importance of objects in the perception of scenes has been a source of behavioral research for decades (Biederman et al., 1982; Davenport & Potter, 2004). Despite this, high-level scene and object visual perception are typically discussed as independent processes, especially with respect to their underlying neural mechanisms. In fact, object-selective and scene-selective brain regions are typically functionally localized by comparing them against one another (Dilks et al., 2013; Epstein & Kanwisher, 1998; Ganaden et al., 2013).

To address this inconsistency, we investigated whether objects within a scene can predict the responses associated with scene perception in three brain regions: the parahippocampal place area (PPA), the retrosplenial complex (RSC), and the occipital place area (OPA); for a review of scene-selective regions see Epstein and Baker (2019). Although considered “scene-selective”, studies have suggested that object properties or the process of object recognition are also reflected in the signal of the OPA, PPA, and RSC. However, in the case of object properties, stimuli used are of individual objects with no background, or a focal object with minimal background (e.g., Mullally & Maguire, 2011; Troiani et al., 2014). In the case of object recognition, stimuli have been either artificially generated (Harel et al. 2013), or manipulated and obscured (Brandman & Peelen, 2017, 2019; Linsley & MacEvoy, 2015). Moreover, the typical inference is that object properties are recruiting scene-selective regions because they help define the spatial nature of scenes (Epstein & Baker, 2019). The field has gained valuable insights from these aforementioned studies, however, to further our understanding we investigated the role that objects play in the neural representation of scenes beyond contributing to spatially defining properties. Critically, we examined how objects modulate the scene-selective signal within a full, unmanipulated, scene with embedded clusters of objects in order to reflect naturalistic visual perception.

Scene categories are often identified by recognizing the objects within them. An important test by Stansbury et al. (2013) found that models built from co-occurring object statistics, representing learned scene categories, were successful at explaining activation patterns in scene-selective cortex. However, their analysis did not isolate the role of objects, but rather suggested that objects are good indices for representing scene categories. We took this a step further and, in our investigation, examined how the variance of objects present within different exemplars of the *same* scene category modulated the pattern of scene-selective signals. By looking within a scene category, we investigated the unique contribution of objects while keeping other properties inherent to the category the same.

To broaden the scope of our results, we studied a diverse set of scenes and objects, and therefore, our sampling of objects, scene categories, and individual scene exemplars are increased by an order of magnitude from the standard study. Typically, only a few scene categories and a handful of objects are varied to examine how scenes and objects interact. This may be prone to confounds related to particular categories and/or objects, and may not provide a realistic assessment of the richness of human scene perception. We used the BOLD5000 dataset (Chang et al., 2019) which includes the fMRI response to viewing almost 5,000 unique scenes. In the current study, we examined 1,179 scenes, comprising 18 different scene categories. The stimuli were color photographs depicting scenes with a number of interacting objects. Each object in a scene was labeled, and then compared across exemplars of the same category (mean = 58 objects/category), as well as across all categories. Using these stimuli, we asked whether scene-preferring regions are sensitive to the objects that comprise the scenes (e.g., a living room with or without a TV, or a bathroom with or without a tub). If so, this would suggest that scene regions are doing more than just processing the unique spatial properties of scenes, but rather processing its contextual makeup (see, Aminoff & Tarr, 2015). For comparison, we looked at an object-preferring region, the lateral occipital complex (LOC) with the expectation that object presence would modulate representations in this region. Together, the results suggest that object presence in scenes affect the neural representation of both scene- and object-preferring regions.

## Materials and Methods

### Dataset

The data used for this study is from BOLD5000 (BOLD5000.org), release 2.0 (Chang et al., 2019, 2021). We used three out of the four participants (the fourth participant only had a portion of the trials). Each participant viewed 4,956 unique scenes over the course of 16 MRI sessions. Although there are few participants, having a within participant dataset across 4,956 unique scenes provides an unparalleled number of unique trials per individual. The participant’s task was to rate whether they liked, disliked, or found the image neutral.

fMRI Data was acquired on a 3T Siemens Verio MR scanner at the Carnegie Mellon University campus using a 32-channel phased array head coil. Functional imaging voxel size was 2^3^ mm. Data was minimally preprocessed, and included motion correction and distortion correction using fMRIPrep 1.1.4 (Esteban et al., 2019; https://github.com/poldracklab/fmriprep). Functional data were analyzed using a customized hemodynamic response function for each voxel, which were then used in a cross-validated general linear model in order to derive a set of optimized noise regressors from voxels unrelated to the experimental paradigm. Beta estimates were then regularized on a voxel-by-voxel basis using ridge regression. Beta estimates for each voxel were then extracted for each region of interest (ROI) for each unique scene. Ten ROIs were used: early visual cortex (EV), lateral occipital complex (LOC), parahippocampal place area (PPA), occipital place area (OPA), and retrosplenial complex (RSC); for both the left (LH) and right hemisphere (RH). Scene-preferring regions (PPA, OPA, RSC) were defined using the contrast of scenes versus weak contextual objects and Fourier transformed scrambled scenes. The LOC was defined using the contrast of weak contextual objects versus Fourier transformed scrambled scenes. The EV regions were defined by the Fourier Transformed scrambled scenes versus baseline, confined to the occipital cortex. See Chang et al. (2019, 2021) for details about fMRI acquisition, data analysis, and ROI definition and extraction.

### Experimental Design and Statistical Analyses

#### Stimuli

Pictures used in the dataset were all color, real-world photographs taken from the ImageNet, COCO, and SUN-like image databases (Chang et al., 2019). In the current analysis, each picture was assigned a basic level scene category (e.g., kitchen, office, street, garden), see Fig 1. Because we wanted to examine how objects might influence the neural representation of a scene, we examined this both across scene categories, and critically, within each scene category. Examining within a scene category reduced confounding the influence of object differences with that of scene category differences, since objects cluster in specific scene categories. To have a sufficient number of trials, we included scene categories that had at least 30 unique exemplars. This resulted in 1,179 stimuli across 18 different scene categories. The median number of unique exemplars per category was 58 (standard deviation 27.6); with a minimum of 34 in train platform, and a maximum of 139 in the street category. For details of number of exemplars per condition, see Fig. 1 caption.

**Figure 1:**
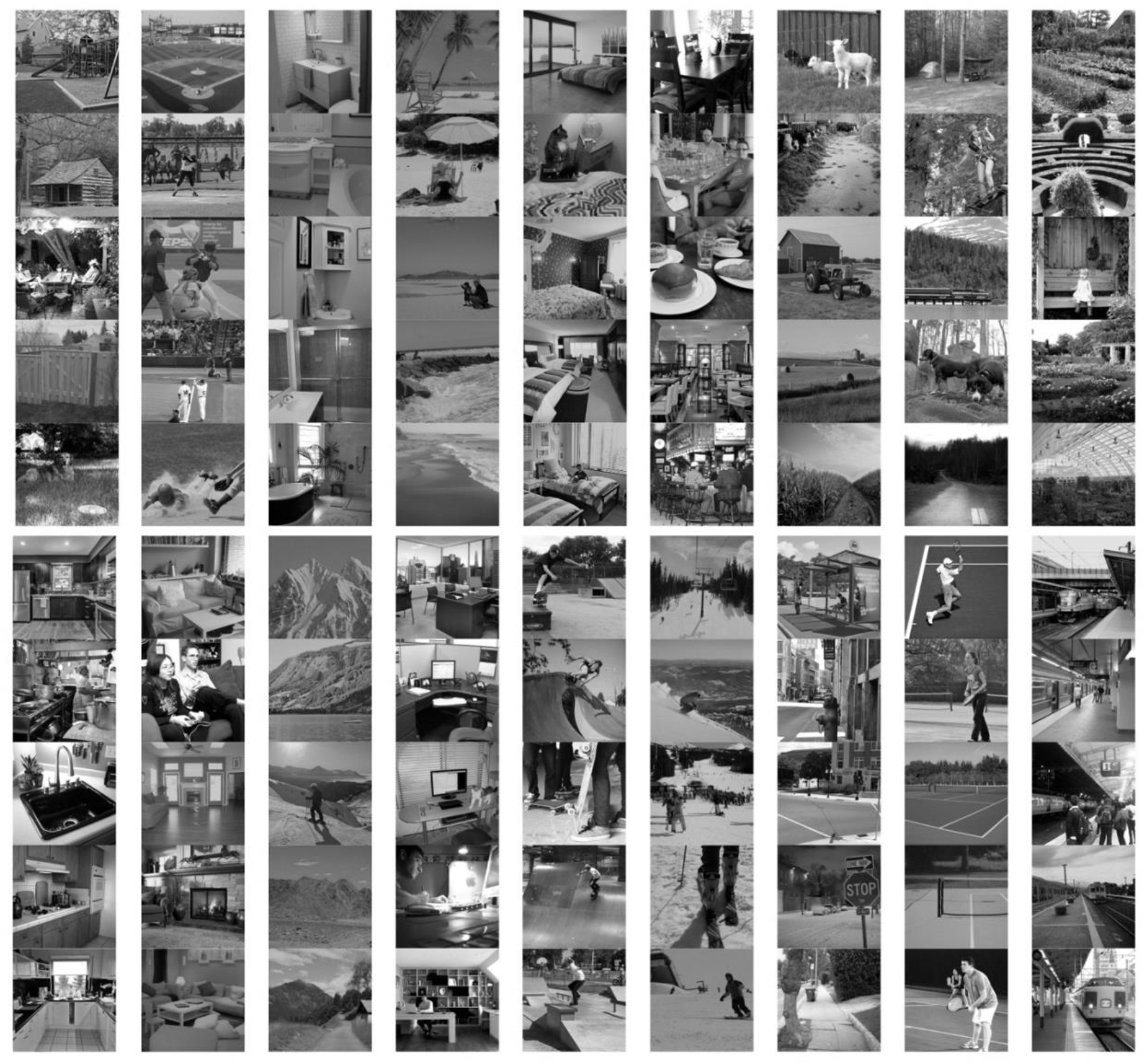
Examples of stimuli. For illustration purposes, pictures are black and white; however, participants in the MRI viewed full color photographs. A total of 1,179 stimuli were divided across 18 scene categories (# of exemplars per category): Backyard (41); Baseball Field (72); Bathroom (88); Beach (65); Bedroom (52); Dining Area (95); Farm (61); Forest (55); Garden (45); Kitchen (83); Living Room (68); Mountain (42); Office (49); Skate Park (35); Ski Slope (104); Street (139); Tennis Court (55); Train Platform (34).

### Object Coding

Object presence was categorized and cross-checked by two lab members and then a third time by the first author, all with the goal of naming as many entities in the scene. A matrix with a binary absence/presence characterization was created detailing what objects were present in each scene exemplar. The object had to be easily recognizable without being too far in the distance and/or too obstructed. Any object that was present in 3 stimuli or less across the entire dataset was coded in a column of misc. large objects and misc. medium/small objects rather than individuating that singular object. This resulted in 266 objects coded across the 1,179 scene stimuli. Only 63 of the 266 object categories appeared exclusively in one scene category, the remaining objects appeared in exemplars in more than one scene category. The median number of scene categories an object was found in an exemplar was three.

#### Data Analysis

Five representational dissimilarity matrices (RDM) were created to understand the representation of scenes in the brain (details below). (1) To model the effect of object variability (e.g., presence of a chair) an object space was generated which compared scenes based on what objects were present in the image using the coded objects described above. (2) To assess low level visual similarity a pixel space was generated to compare each scene based on raw pixel similarity and (3) a gabor wavelet similarity space was generated. (4) To model the effect of semantic scene category (e.g., kitchen vs garden) a category similarity space was generated. Lastly, (5) an fMRI space was generated from comparing the pattern of fMRI BOLD activity across voxels in different regions of interest (ROI) in the brain for each scene. The goal was to see if object presence, above pixel and gabor similarities, could explain the similarities and dissimilarities in the fMRI space, which would reveal whether the nature of scene representation included objects in each ROI.

*Object Space* was created through a matrix of the objects present across all scene exemplars. Each exemplar was then represented by a vector of 1s and 0s denoting which objects were present in that specific picture. A principal components analysis was then performed to reduce the number of features in the matrix. The components that accounted up to 70% of the variance were included for further analysis. An RDM was generated by computing the Euclidean distance between pairwise comparisons of the principal components.

*Pixel Space* for the scenes was generated using the sum squared error (SSE) pixel tool in the Image Similarity Toolbox (Seibert, 2013). This method sums the squared difference of pixel values for each image. One image is linearly shifted in the x and y directions over the image being compared. The shift which results in the lowest SSE is kept. Each pairwise comparison was analyzed for pixel differences to create a pixel space RDM. These similarity scores were then z scored for the final pixel space RDM.

*Gabor space* for the scenes was generated using the Gabor filterbank tool in the Image Similarity Toolbox (Seibert, 2013). This method projects the image onto a Gabor wavelet pyramid as a model for primary visual cortex. The filters span eight orientations (multiples of .125*pi), four sizes (with the central edge covering 100%, 33%, 11%, and 3.7% of the image), and X,Y positions across the image (such that filters tile the space for each filter size). The resulting vector of filter responses are compared between images using the Euclidean distance to create an RDM.

*Category space* for the scenes compared the semantic category across each pairwise comparison of scenes. Thus, each scene was assigned one of the 18 basic level categories, and for each pairwise comparison it was coded either as the scenes belonged to two different categories, or the same category (regardless of category). Thus, only two codes were used: different (−1) and same (1).

*fMRI Space* was generated by creating an RDM from each ROI data from the BOLD5000 dataset. Ten ROIs were used: early visual cortex (EV), lateral occipital complex (LOC), parahippocampal place area (PPA), occipital place area (OPA), and retrosplenial complex (RSC); for both the left (LH) and right hemisphere (RH). For each ROI, a matrix of scene exemplar by voxels were used, with each cell containing the beta weight extracted for that particular voxel, for a specific scene. A principal components analysis was then performed to reduce the number of features. Those components that accounted up to 70% of the fMRI activity were included. To generate the RDM, we calculated the Euclidean distance across each pairwise comparison of scenes to create an RDM for each of the three participants. These individual participant RDMs for each ROI were then averaged across participants to create a single RDM for each ROI, which was used in the analysis on the average. All analyses were performed both on the average RDM and on each individual RDM to see the consistency of the effect. The fMRI space was used as the neural representation.

##### Regression Analysis

We used a linear regression to determine whether variance in the object composition of a scene was reflected in the neural representation of a scene. A linear regression was performed on each of the ten ROIs. Each ROI was the dependent measure to predict. A hierarchical model was used in which pixel space, gabor space, and category space (if applicable) was first entered into the model to determine how much of the fMRI similarity space could be accounted for by low level visual properties and by semantic category. We then entered the object RDM to determine whether there was a significant amount of variance accounted for by the object RDM over and above the pixel, gabor, and semantic category RDM. This was reflected in the change in r with the addition of the object model. Unstandardized betas and associated standardized error were extracted from the final model. Variables were assessed for collinearity, with no significances found. If the change in r was significant with the addition of the object model, this would support that objects are incorporated into the neural representation of a scene for the examined ROI.

Three different regression analyses were performed. The first, detailed in the paragraph prior, was a regression predicting the pairwise similarity of all 1179 scene stimuli, resulting in 694,431 pairwise comparisons values for each RDM. This analysis is referred to as the All scenes (All) analysis. A second analysis removed those pairwise comparisons that included scenes of the same semantic category. This resulted in looking at the effects of object similarity embedded within scenes across different categories, referred to as different scenes analysis (Diff). The RDMs used in the analysis included 649,955 pairwise comparisons. The final analysis looked only at the comparisons of scenes within the same category, referred to as Same category scenes analysis (Same). For this analysis, matrices were split into individual matrices for each category. Thus, only those objects which were present across the exemplars of that particular category were included in the object matrix. A principal components analysis was then performed on each of the 18 category object matrices, and a Euclidean distance RDM generated from those components that accounted for up to 70% of the data. Similarly, the fMRI data was separated into matrices for each separate category. A principal components analysis was then performed on these category fMRI ROI matrices, and a Euclidean distance RDM generated from those components that accounted for up to 70% of the data. Pixel and Gabor matrices were generated for just within category. The resulting category specific matrices were then concatenated to one large matrix. The resulting matrix included 44,476 pairwise comparisons entered into the regression. In the regression for the Diff and Same analyses, the category space model was not entered since it was no longer relevant.

##### Significance testing

Significance of the object model as evidenced by the change in r^2^ was evaluated in a bootstrapping method. Separately for each analysis (All, Diff, and Same, and for each individual participant), the object model was resampled randomly with replacement. For each resampled object model, the same regression analyses as described above were run, resulting in a change in r^2^ related to the resampled data. To generate a confidence interval, the data was resampled 1000 times, to provide 1000 r^2^ data points. Significance of p < .05 can be determined as the r^2^ generated from the real data does not fall within 95% of the distribution of the resampled r values. If the r^2^ generated from the real data is completely non-overlapping with the bootstrapped data, we can assume p < .0001.

To compare the effect of object model across different ROIs and across the Diff and Same analyses unstandardized betas were used. Unstandardized betas and standard error for the object model were extracted from the final model of the regression. For a contrast of interest, this equation from Paternoster et al. was used (Paternoster et al., 1998).

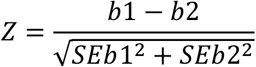

The Z score was then used to assess the p value for the contrast of interest. The p value in these analyses were Bonferroni corrected for multiple comparisons. When comparing across regions, the correction was for 36 comparisons (12 for each analysis x 3 analyses per individual); when comparing across the Diff and Same analysis the correction was for 30 comparisons (10 ROIs across x 3 analysis).

## Results

We examined whether the presence of specific objects in a scene would be reflected in the neural representation of a scene, and in particular, in scene-selective brain regions. If the object model predicts the neural response in scene-selective regions, this would suggest that objects are processed by the same neural mechanisms, and are thus a component of, scene perception. To address this, a linear regression was used to assess how differences in objects present in each image could account for differences in the neural representation of scenes across ten different regions of interest. Pixel, gabor, and category similarity were first entered into the regression to isolate the effect related to objects and to avoid confounds such as low level visual similarity and category similarity. Then, the object model was entered to test if the object model could account for variance over and above that of these low-level visual models. Significance was assessed with a bootstrapping confidence interval, and with a Bonferroni correction for multiple comparisons.

For the first analysis, we used all 1,179 scenes, which included scenes from the same semantic category, and scenes from different categories. The object model was significant in all three participants in the LH EV, LH PPA, and the RH PPA at p < .0001. The model was additionally significant across all three participants with p < .05 (not all withstanding correction for multiple comparisons) in RH EV, RH LOC, RH OPA and RH RSC. In addition to looking at the reliability of the effect across the individual participants, we also created an averaged fMRI similarity space to increase the signal to noise in the RDM. When using the averaged fMRI similarity space, the object model in the All analysis was significant p < .0001 in all ROIs except for the RHEV.

Next, we asked whether scenes that are in different categories (e.g., a living room and an office) were treated as more similar if they had similar objects (e.g., couch). This analysis (Diff), which only included only the comparisons of those scenes that differed in semantic category. In this analysis, we found the object model was significant across all three participants at a p < .0001, in the LH EV, bilateral PPA, RH LOC, and RH RSC. Using a less stringent p < .05, the object model was also significant across all three participants in LH LOC. When using the average data, the object model was significant in each ROI at p < .0001, except for RH EV which was only significant at p < .05, see Fig. 2 and Fig. 3.

**Figure 2:**
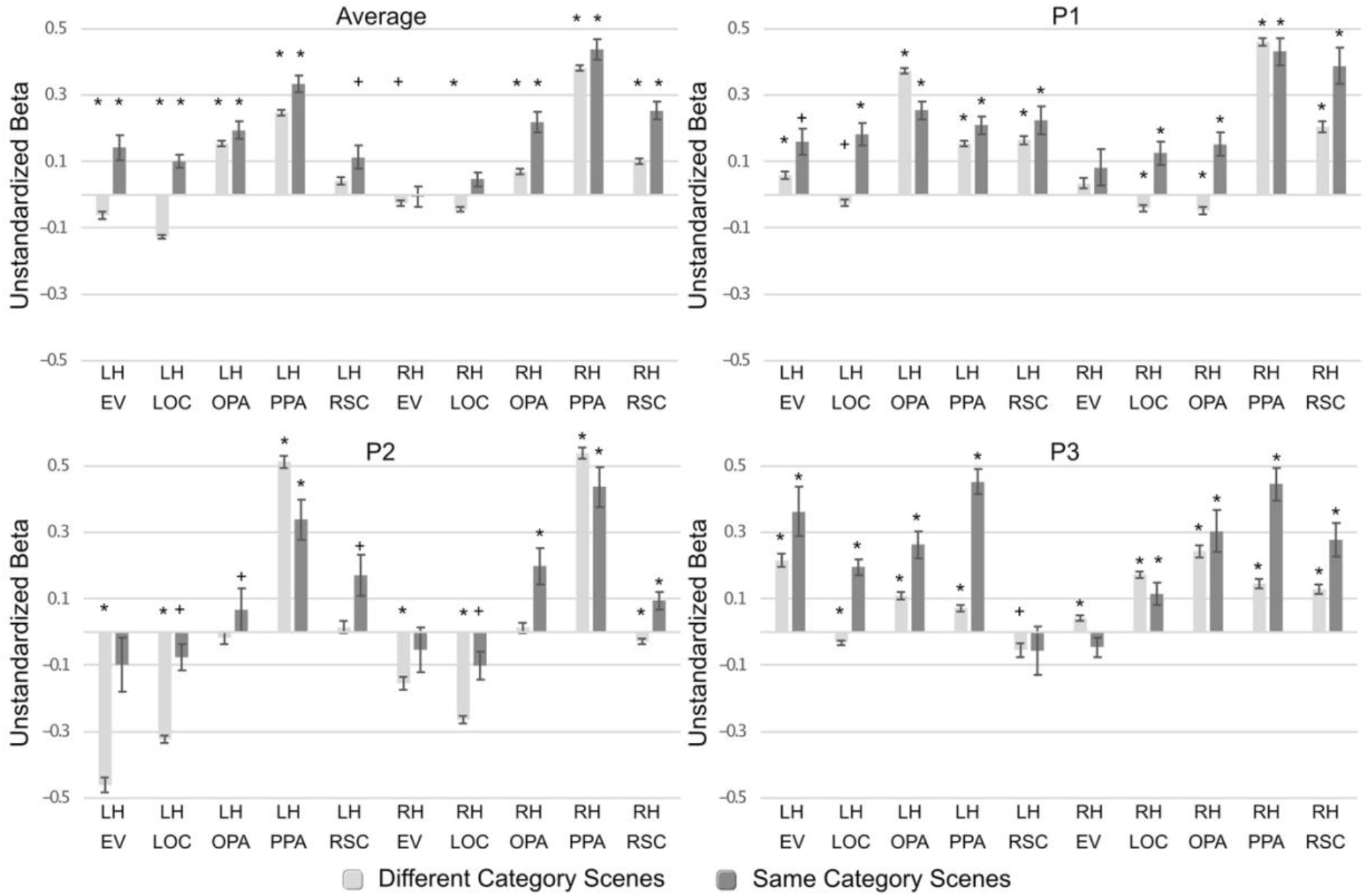
Unstandardized Beta scores of the object model for the three participants (P1, P2, and P3) and the average. Analysis comparing scenes across different categories in light gray; analysis comparing scenes within the same categories displayed in dark gray. Significance of the object model is indicated with * for p < .0001 and + p < .05. Significance is based on the bootstrapping confidence interval tests on the change in r^2^ when adding the object model in the regression over and above pixel and Gabor similarity. Error bars are standard error derived from the final model of the regression.

**Figure 3.**
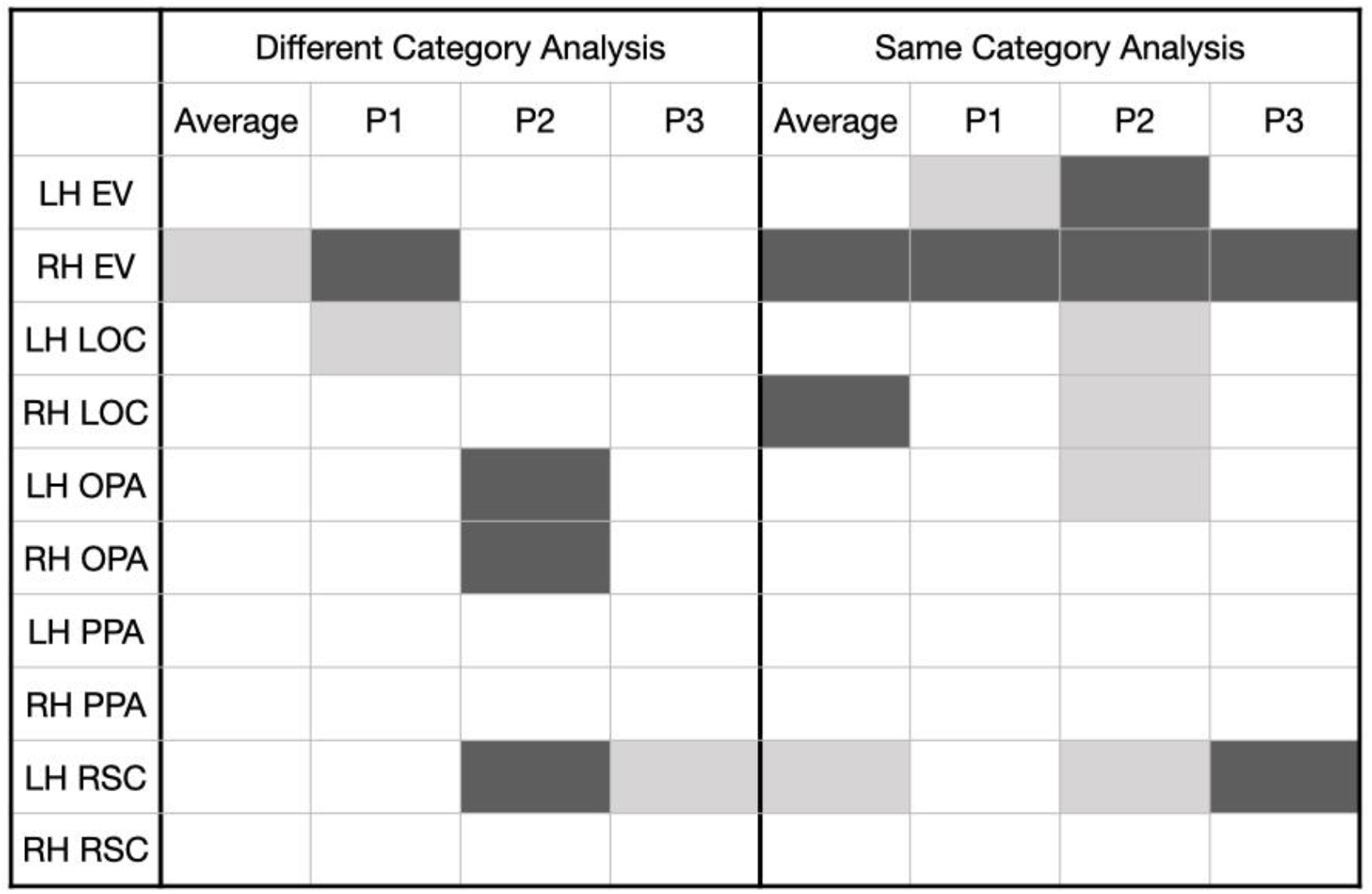
Visualization of the significant ROIs in the Diff category analysis and the Same category analysis. White (no fill) cells indicate the object model accounted for a significant amount variance over and above the pixel and gabor model correcting for multiple comparisons. Light gray shaded regions indicate significant p < .05, uncorrected. Dark gray shaded regions indicate no significance.

We then asked whether these effects would also be significant in comparisons of scenes within the same category, where you might expect the objects of a scene to have more weight to differentiate between exemplars of the same category (e.g., a living room with and without a TV). In this analysis (Same), we found the object model to be significant across all three participants at a p <.0001 in bilateral PPA, RHOPA, and RHRSC. When dropping the significance down to p < .05, the object model was also significant across all three participants in the bilateral LOC, and LH OPA. When using the averaged data, the object model was significant at p < .0001 in LH EV, LH LOC, bilateral OPA, bilateral PPA, RHRSC, and at p < .05 in LH RSC.

From this analysis, we can determine that objects play a significant role in the representation of scenes, especially in scene-selective regions. Using the z scores obtained from assessing the difference between unstandardized beta values we assessed whether the object model was more significant in scene-selective ROIs compared with an object-selective region, and whether there were significant differences across each of the three scene-selective regions. These comparisons were performed only within hemisphere and only used the Diff analysis and the Same analysis, since these together comprise the All analysis. When comparing scene-selective regions to the LOC in the Diff analysis, the object model was significantly greater p < 0.0016 (which is a p < .05 with Bonferroni correction for 32 comparisons) across all three participants in the LH OPA and LH PPA compared to the LH LOC, no significant differences were consistently found in the RH withstanding correction for multiple comparisons. Using the averaged data, which may be more stable, significant difference were found when comparing all scene regions with LOC in both hemispheres, with stronger role of objects in the scene regions compared with the LOC (see Table 1, top left section).

**Table 1.** Z score results from beta comparisons across difference regions and analyses. Bold values highlight when all three participants had significant effects. Standard text indicates significant with correcting for multiple comparisons. Gray values indicate significance at p < .05 without correcting for multiple comparisons. Shaded values indicate no significance. Comparisons of scene-preferring regions compared with LOC in the top section; comparisons across scene-preferring regions in the second section; hemispheric comparisons in the third section; and in the last section, comparisons across the analysis with using scenes that differed in semantic category (Diff) compared to the analysis using only scenes within the same category (Same).

| Contrast | Different Categories Analysis |  |  |  | Same Categories Analysis |  |  |  |
| --- | --- | --- | --- | --- | --- | --- | --- | --- |
|  | Average | P1 | P2 | P3 | Average | P1 | P2 | P3 |
| LH OPA vs. LOC | **28.70** | **31.71** | **13.71** | **10.84** | 2.77 | 1.67 | 1.86 | 1.46 |
| RH OPA vs. LOC | 10.38 | -0.52 | 14.00 | 3.36 | 4.58 | 0.54 | 4.40 | 2.64 |
| LH PPA vs. LOC | **38.71** | **14.75** | **39.03** | **8.19** | 7.21 | 0.61 | 5.84 | 5.83 |
| RH PPA vs. LOC | 38.61 | 31.49 | 37.51 | -1.54 | **10.50** | **5.58** | **7.36** | **5.48** |
| LH RSC vs. LOC | 14.25 | 12.13 | 15.95 | -0.98 | 0.30 | 0.77 | 3.40 | -3.31 |
| RH RSC vs. LOC | 14.25 | 12.74 | 16.59 | -2.50 | 5.95 | 4.00 | 3.99 | 2.65 |
| LH OPA vs PPA | -8.14 | 19.72 | -20.13 | 2.45 | -3.66 | 1.20 | -3.05 | -3.47 |
| RH OPA vs PPA | -24.65 | -31.91 | -22.05 | 4.16 | -5.11 | -5.06 | -2.94 | -1.77 |
| LH OPA vs RSC | 8.63 | 14.07 | -1.24 | 6.75 | 1.81 | 0.60 | -1.16 | 3.85 |
| RH OPA vs RSC | -2.67 | -13.12 | 2.35 | 4.84 | -0.86 | -3.57 | 1.72 | 0.32 |
| LH PPA vs RSC | 15.74 | -0.72 | 19.58 | 5.23 | 5.02 | -0.31 | 1.94 | 6.22 |
| RH PPA vs RSC | 23.27 | 12.58 | 29.13 | 0.87 | 4.54 | 0.62 | 5.19 | 2.35 |
| LH vs. RH EV | -2.58 | 1.32 | -10.45 | 8.04 | 2.97 | 1.17 | -0.43 | 5.04 |
| LH vs. RH LOC | -9.80 | 1.15 | -3.57 | -17.40 | 1.88 | 1.19 | 0.46 | 1.93 |
| LH vs. RH OPA | 7.13 | 32.16 | -1.23 | -6.17 | -0.57 | 2.30 | -1.53 | -0.56 |
| LH vs RH PPA | -11.34 | -21.37 | -1.03 | -4.18 | -2.62 | -4.46 | -1.16 | 0.11 |
| LH vs RH RSC | -4.56 | -1.95 | 2.23 | -6.98 | -3.15 | -2.34 | 1.16 | -3.78 |
| Same vs. Different Comparison |  |  |  |  |  |  |  |  |
| LH EV | 5.04 | 2.50 | 4.29 | 1.88 |  |  |  |  |
| RH EV | 0.65 | 0.86 | 1.44 | -2.79 |  |  |  |  |
| LH LOC | **11.38** | **5.94** | **6.15** | **9.51** |  |  |  |  |
| RH LOC | 3.98 | 4.47 | 3.82 | -1.62 |  |  |  |  |
| LH OPA | 1.38 | -4.25 | 1.23 | 3.68 |  |  |  |  |
| RH OPA | 4.69 | 5.35 | 3.28 | 0.94 |  |  |  |  |
| LH PPA | 3.23 | 1.99 | -2.81 | 9.72 |  |  |  |  |
| RH PPA | 1.77 | -0.67 | -1.62 | 5.81 |  |  |  |  |
| LH RSC | 1.95 | 1.38 | 2.45 | -0.04 |  |  |  |  |
| RH RSC | **5.39** | **3.19** | **4.43** | **2.82** |  |  |  |  |

When looking only at those comparisons of scenes within the Same category, the only significant difference with correction across all three participants was found with the object model having a significantly greater beta in the RH PPA compared with the RH LOC. Using the averaged data, the object model had a stronger role in the LH PPA vs LH LOC, and RH OPA, RH PPA, and RH RSC compared with RH LOC all at a significance of p < .0016 (see Table 1, top right section). The next set of comparisons were performed to see if the object model was significantly different in any of the three scene-selective brain regions across all three participants (Table 1, middle second section). There was no significance found comparing across the three brain regions in all three participants, in the same direction, with correcting for multiple comparisons. When using the averaged data, in the Diff analysis, an object model was stronger in the PPA than the OPA in both hemispheres, PPA than RSC in both hemispheres, and in OPA compared with RSC in the LH. For the Same analysis, the object model was stronger in PPA than OPA or the RSC in both hemispheres, see Table 1.

The data seemed to trend with a more significant effect of the object model in the right hemisphere compared to the left. However, we did not find this to be a consistent effect across our participants (see Table 1, third section). Using the averaged data, we did find a significant difference in the Diff analysis in the LOC, OPA, PPA, RSC; and in the Same analysis in the EV and the RSC. Moreover, there was a trend for the object model to have a higher beta in the analysis with just the comparisons of scenes within the same category compared to the comparisons of scenes across categories. This did have a significant effect across all three participants in the LH LOC and the RH RSC where the object model had a significantly stronger beta in the Same analysis compared with the Diff analysis across all three participants (Table 1, bottom section). When analyzing the average data, significance was also found for this contrast in the LH EV, bilateral LOC, LH PPA, RH OPA, RH RSC. For a complete set of Z scores for all comparisons, please see Table 1.

## Discussion

This study explored whether objects are integrated in the neural representation of scenes. We examined how the presence of objects across different exemplars of the same scene category (e.g., a garden with a bench vs. a garden with a pond) and across different categories (e.g., a TV in living room and a TV in a bedroom) predicted the representation of scenes in scene-preferring regions: OPA, PPA, and RSC in comparison to an object-preferring region, LOC. Since objects cluster in particular scene categories, it was critical to examine this both *within* a scene category to ensure that object presence was not confounded with other features of scene categories (Greene & Oliva, 2009; Oliva & Torralba, 2006) and across categories. Our results suggest that: 1) The variation in objects embedded within a scene predicted the scene’s neural representation in both object-preferring **and** scene-preferring regions. 2) The variation of objects within scenes had a stronger role when comparing the representations of scenes within the same category compared to across categories. 3) When comparing scene-preferring regions with object-preferring regions, if significant differences were found, it was for the object model to modulate the representations in scene-preferring regions more than object-preferring regions.

The results stress the function of “scene” processing and “scene-selective” brain regions should be thought of as processing scenes in conjunction with the embedded objects rather than in their absence. Critically, we provide direct neural evidence that scenes are not processed or represented in isolation from their constituent objects. This is relevant not only when objects may indicate scene category (e.g., Stansbury et al., 2013), but also when distinguishing between different exemplars of the same scene category, and when scenes share the same objects, but of different semantic categories.

We traditionally associate the processing of objects with the LOC. The LOC is sensitive to shape, object category, viewpoint invariance, and interacting objects (Grill-Spector et al., 1999, 2001; Kim & Biederman, 2011; Kourtzi, 2001; Malach et al., 1995; Sawamura, 2005). Our results suggest that the preferred stimulus of the LOC may not be objects and shapes at large. If it were, we would expect it to have a stronger relationship to an object model than scene-preferring regions, which we did not find. When comparing the degree to which the object model accounted for variance in the response of the scene-preferring regions to the LOC, we found that either there was no significant difference, or when there was, it was for there to be a greater effect in the scene-preferring regions than LOC (e.g., LH OPA, bilateral PPA). This may support the finding that the LOC is sensitive to shape interacting with retinotopic or salient positioning (Gomez et al., 2019; Levy et al., 2001; Sayres & Grill-Spector, 2008): i.e., the LOC is most sensitive to single objects that are salient in our central vision. In our analysis of complex, real-world scenes, we hope to add to the literature about how to understand the functional role of the LOC in object processing. Future research will need to determine whether the similarity between an object model and LOC activity would increase, and be significantly greater than scene-selective regions, if only modeling the object that is foremost in the scene. However, the results of the current study emphasize the integrative nature of object and scene understanding. The results provide support for the role of context in object perception (e.g., Cacciamani et al., 2014; Flowers et al., 2020; Quek & Peelen, n.d.; Skocypec & Peterson, 2022) and that scene processing inherently includes the processing of the associations embedded within it (Aminoff, 2014; Aminoff & Tarr, 2015, 2020).

There were subtle differences between scene-preferring regions. Bilateral PPA was the most consistent region modulated by the object model – this was significant across all three participants individually and in the average. The object model also consistently accounted for variance in the RH RSC – significant across all three participants individually and in the average. Numerically, there was a stronger effect in the right hemisphere compared with the left hemisphere. Although this was not found to be statistically significant across all three participants, it was in the same direction in all three participants, and statistically significant in two out of the three participants in LOC and PPA in the comparisons of scenes across different categories. Interestingly, this effect largely disappeared when comparing scenes within the same categories. This may relate to evidence suggesting that the RH is more sensitive to the details of a scene. For example, the RH PPA has been associated with visual properties of scenes, whereas the LH PPA with semantic-conceptual properties of scenes (Stevens et al., 2012). Topographic agnosia and disorientation has also been associated with RH deficits (Aguirre, 1999). Bilalić et al. (2019) found the RH PPA to be sensitive to the object composition of a scene. This may suggest the RH scene representation is sensitive to the details that differentiate between scenes, and recognizing when scenes have commonalities (i.e., objects) that cross semantic categories.

There were also differences found when comparing the analysis which included comparisons only from scenes from different categories to the analysis which included comparisons only from scenes in the same category. Namely, the object model played a stronger role in the Same analysis than the Diff analysis. Note, there was a significant difference in the number of comparisons in each analysis with the Diff analysis having considerably more. However, if the differences were a reflection of the number of comparisons, then we would expect to have noisier results with the Same analysis, but we found consistent results showing a stronger effect in the Same analysis. Moreover, the strength of the object model predicting similarity across representations of scenes within the same semantic category should better isolate the role of objects, because other co-occurring statistics that may vary across scenes from different semantic categories were held constant.

The stronger effect of the object model in the Same analysis was significant across all three participants in the LH LOC and the RH RSC. And in the same direction, and significant across two participants in the RH LOC and the RH PPA. This makes intuitive sense in that objects help in distinguishing between exemplars of the same category such as a living room with and without a TV, or a bedroom with bunk bed versus a crib. Objects may give rise to different affordances within a scene category, and affordances have proven to be an important aspect of scene understanding and representation (Auger et al., 2012; Greene et al., 2016; Janzen & van Turennout, 2004). More work will need to be done to examine why the LOC demonstrated such significant differences between the two analyses. For example, does distinguishing between exemplars of the same category engage the type of object processing more associated with the LOC? Moreover, the RH RSC showing a difference demonstrates a contribution of the RH RSC in understanding the composition of a scene, especially with regard to the semantic components. These differences in the results, yielded from the analysis of comparing scenes across different semantic categories and scenes within the same semantic category highlights the importance of looking at scene processing in each of these domains. Different features may be more relevant when examining scenes of the same category compared with scenes of different categories, and looking at both types or processing provides a more complete understanding of scene perception.

Neuroimaging research has used functional localizers to define scene-selective regions and object-selective regions. Typically, in these localizers, single objects are presented front and center without background, or with non-naturalistic background, and are compared to scenes. Voxels more active for objects are ascribed to object-selective regions and voxels more active for scenes are ascribed to scene-selective regions. Our data proposes these regions are not exclusive for one stimulus type compared to another, and thus the contrast employed may be missing relevant voxels. A limitation of our study is that the effects were small. We are not claiming that objects can account for all of a scene representation. There are many important features of a scene not captured by an objects analysis, such as navigational affordances, affordances more generally, three-dimensionality, textures, colors, peripheral visual field differences, geometrical relations, contextual associations more general in both spatial and non-spatial domains, etc. The purpose of the study was to investigate whether objects embedded within a scene are included in the scene representation rather than processed exclusively in the LOC, known to be involved in object perception.

In conclusion, the functional role of scene-preferring regions must not be thought of as scene-only regions at the exclusion of objects, but rather, should include objects in the representation of these regions. Objects are integral to the processing and understanding of scenes. Visual processing regions may be functionally defined not as a response to one stimulus category versus another, but rather with respect to which processes are engaged when interacting with the stimulus category, such as processing groups of objects in scenes, or processing a single salient object in our foreground.

## Acknowledgments

This work was supported by National Science Foundation (grant number BCS 1439237) and internal support from Fordham University. We thank Kate Uhling, Daphne Baker, Michael L’Abbate, Edona Gjonbalaj, Emma Kreutzmann, Emily Huegler, Benedict Antonyraj, Elizabeth Galbo, and Madison Gadea for their help in labeling objects in each scene. We thank Jacob Prince for his help with regard to the BOLD5000 release 2.0 data.

The authors declare no competing financial interests.

